# Applying Inter-rater Reliability to Improve Consistency in Classifying PhD Career Outcomes

**DOI:** 10.1101/370783

**Authors:** C. Abigail Stayart, Patrick D. Brandt, Abigail M. Brown, Tamara Hutto, Rebekah L. Layton, Kimberly A. Petrie, Emma N. Flores-Kim, Christopher G. Peña, Cynthia N. Fuhrmann, Gabriela C. Monsalve

## Abstract

In the past year, there has been an exciting groundswell of national efforts to integrate multiple taxonomies for the transparent dissemination and analysis of PhD career outcomes. In this study, we leveraged the unique resources of the Broadening Experiences in Scientific Training Consortium to examine the reliability of the three-tiered Unified Career Outcomes Taxonomy (UCOT v.2017) that was collaboratively developed at a meeting convened by Rescuing Biomedical Research in August 2017. Using an amended version of the UCOT v.2017 (UCOT v.2017-rev1) and a new Supplementary Guidance document, we categorized over 570 PhD alumni records from three different universities. Utilizing Krippendorff’s alpha to measure the interrater reliability from nine different individuals, we determined moderate to robust reproducibility within the first two tiers of the taxonomy (Workforce Sector and Career Type); however, the reliability for the third tier (Job Function) did not meet established standards. The team identified significant sources of error, revised category definitions, improved coder training materials and processes, and tested for improved reliability through coding 219 PhD alumni records using the revised taxonomy, UCOT v.2017-rev2. Our results revealed that the changes introduced in UCOT v.2017-rev2 improved inter-rater reliability in all three tiers, and either met or exceeded the acceptable standards for reliability. A final set of clarifications were made to UCOT v.2017-rev2, resulting in UCOT v.2018 and a Finalized Guidance document. Our findings underscore the importance of carefully developing guidance documents to aid coders in the reliable and consistent categorization of alumni career outcomes. We propose periodic assessment of the UCOT v.2018 to address the natural evolution of PhD careers in the global workforce. Ultimately, we hope that UCOT v.2018 will aid in the classification and dissemination of alumni career outcomes that is essential to educating trainees, institutions, and agencies about the diversity of career options for PhDs, and therein empower all PhDs to pursue the careers of their choice.

## INTRODUCTION

Recently, a national conversation regarding the diversity of career trajectories of PhDs highlighted the need for greater institutional transparency in reporting career outcomes of graduate and postdoctoral alumni. In addition to fulfilling institutional reporting requirements, public sharing of outcomes data provides information to prospective and current trainees about the broad career paths that leverage their PhD training. In response to the current lack of career outcome visibility, organizations and funding agencies have made concerted efforts to encourage and support institutional commitment to public sharing of career outcomes data. Many groups have called for institutional transparency in career outcomes in the PhD-trained biologists, including, but not limited to: National Institutes of Health [1], Future of Bioscience Graduate and Postdoctoral Training Conference (FOBGAPT) I & II Conferences [2, 3], Rescuing Biomedical Research (RBR) [4, 5], Future of Research (FoR) [6], UW-Madison Workshop [7], National Institute of Health Broadening Experiences in Scientific Training Consortium (NIH BEST) [8, 9], Coalition for Next Generation Life Science (NGLS Coalition) [10], Council of Graduate Schools (CGS) [11], Association of American Universities (AAU) [12], Association of American Medical Colleges (AAMC) [13], and the National Academy of Sciences [14, 15]. Many institutions are now publicly sharing their alumni career outcomes data on websites and in publications [8, 16, 17, 18, 19]. However, a major impeding factor in this effort to share outcome data has been the multitude of career outcomes taxonomies developed by individual institutions and agencies. The absence of a unified language for reporting the career outcomes of graduate and postdoctoral alumni perpetuates an environment in which universities and prospective trainees are unable to compare career outcomes on a local, regional, or national level.

In Spring 2017, fourteen schools within the BEST Consortium formed a working group to design a taxonomy of career descriptions that could be used across institutions to consistently describe the career outcomes of PhD and postdoctoral alumni trained in the biological and biomedical sciences. The working group used the Science Careers myIDP career categories as its starting point for the taxonomy. This work was subsequently incorporated into a collaborative effort led by RBR, which convened a diverse set of national stakeholders, including representatives from AAMC, the NIH, the AAU, and academic institutions both internal and external to the BEST Consortium. The resultant three-tiered Unified Career Outcomes Taxonomy v.2017 (UCOT v.2017) was first released online by the BEST Consortium [20], and reports of the meeting, including additional recommendations for data collection, were reported by RBR, AAMC, and BEST [5, 13, 21, 22].

The UCOT v.2017 has three levels of classification: Workforce Sector, Career Type, and Job Function. In particular, the Job Function tier provides for a granular refinement of the specific skill sets and/or credentials required for employment within that function. Each taxonomic category of UCOT v.2017 was originally developed through collaborative discussions with experts in career development and biomedical training outcomes, and agreed upon at the August 2017 RBR meeting, with the aim of creating a publicly available, standardized, and valid measure that was vetted by experts at the national level. The process of developing and adopting the UCOT v.2017 and its associated recommendations was an exceptional example of cross-organizational communication, collaboration, and compromise.

While the categories in each tier were deemed sufficiently broad to describe the primary career trajectories of PhDs in the biological sciences, the breadth of those definitions permitted significant room for discordant interpretation. The authors of this study were concerned that the breadth of the definitions would obscure the nuances of the complex taxonomy and result in unreliable and inconsistent application of the taxonomy across different institutions, thereby invalidating cross-site comparisons and diluting the value of national reporting. While “reliability does not guarantee validity…*unreliability* limits the chances of validity” [23]. Without a reliable, reproducible, and robust schema for the classification of career outcomes, the national effort to report these data would be severely hampered. The authors believed it crucial to test the reliability of the Unified Career Outcomes Taxonomy v.2017, uncover any inconsistencies in the coding of jobs according to its taxonomic tiers, and use the results to iteratively refine the taxonomy until all tiers met or exceeded reliability standards. In this study, we identified taxonomic categories that resulted in low concordance among coders (*defined here as inter-rater reliability*), and iteratively modified the taxonomy until all three tiers reached suitable levels of reliability among the raters.

## MATERIALS AND METHODS

Institutional Research Board (IRB) approvals for this project were obtained from each institution that provided alumni records (*Emory University: IRB# H13506; Vanderbilt University School of Medicine: IRB #180315; University of North Carolina, Chapel Hill: IRB # 14-0544*).

### Procedure

The measure chosen as most appropriate to examine categorical consistency across raters was Krippendorff’s Alpha [24], which estimates the level of agreement (*inter-rater reliability*) among raters. Parameters of acceptable reliability using Krippendorff’s alpha start at a lower bound of 0.67 (*for a review, see* [25]*; estimates range from 0.67 - 0.80*), with 0.70 being the convergent recommendation used to define reliability among raters ([23] *gives the caveat that 0.67-0.80 should be interpreted with caution*). Hence, we set 0.70 as our comparison point for an acceptable measure of reliability and replicability of the taxonomies tested in this work. Inter-rater reliability was calculated using SPSS software package (Version 24), with an amended macro designed for Krippendorff’s alpha [26].

Preliminary application of UCOT v.2017 (*not reported here*) by four coders from the BEST Consortium (*PDB, TH, AMB, CAS*) revealed several areas of confusion. Therefore, prior to beginning Round 1 of the experiment reported here, this group agreed upon initial revisions to the taxonomy UCOT v.2017 (generating UCOT v.2017-rev1) and two of the group members (*TH and CAS*) developed a draft Guidance Document to specifically address topics or categories that were anticipated to result in discordant interpretation. The group of raters involved in Round 1 of the experiment included 4 experienced coders *(involved in experimental design and initial revisions to the UCOT v.2017)* and 2 naïve coders recruited to join the experiment (**Figure 1**).

**Figure 1:**
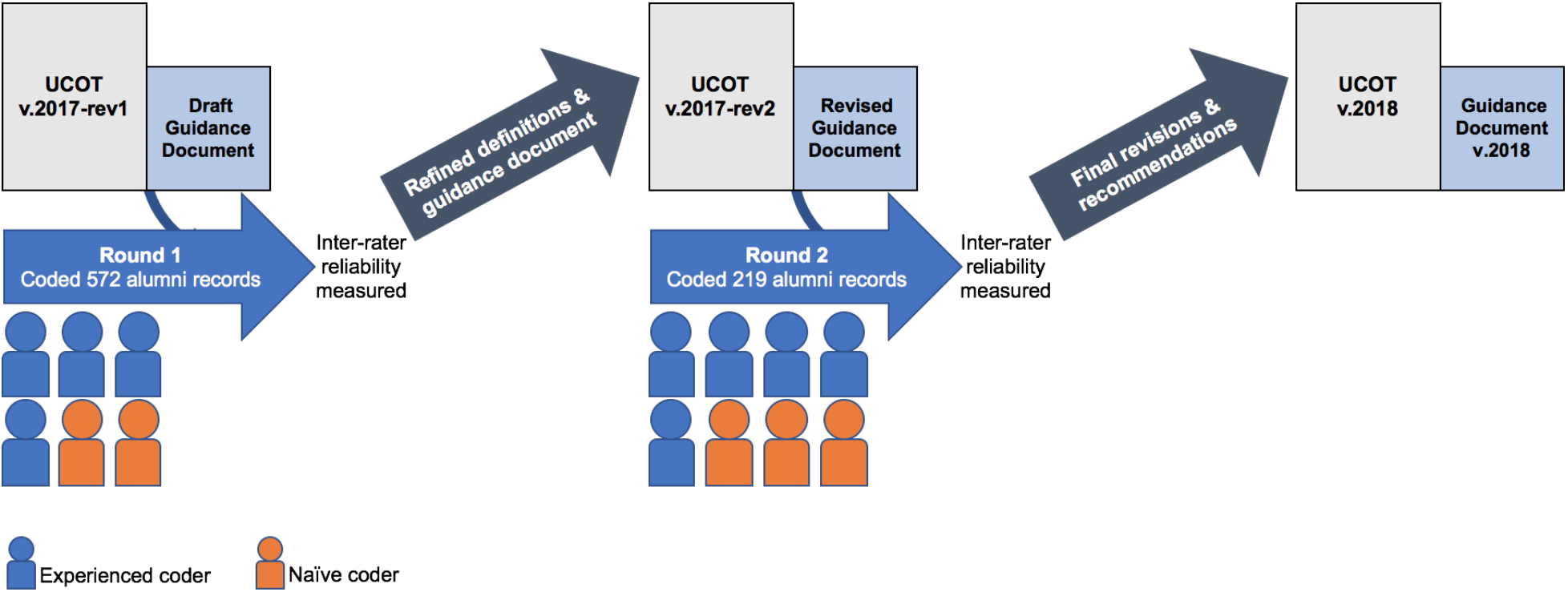
Experimental Design to Assess Inter-Rater Reliability of the Unified Career Outcomes Taxonomy. In Round 1, six raters— comprising four experienced coders *(blue)* and two naïve coders *(orange)*— classified 572 PhD alumni records from three different institutions using UCOT v.2017-rev1 and the draft Guidance Document. After measurements of inter-rater reliability were determined, the working group convened to refine the definitions and guidance materials, generating the UCOT v.2017-rev2 and Revised Guidance Document. The reliability of UCOT v.2017-rev2 and the Revised Guidance Document were then assessed using a new data set of 219 PhD alumni records. In Round 2, coding was performed by five experienced coders (*blue*) and three new naïve coders (*orange*). After categorization, inter-rater reliability was measured. The group convened again to review the results and agree upon final recommendations for revisions to the taxonomy and guidance document, thereby generating UCOT v.2018 and Guidance Document v.2018.

In Round 1 of the experiment, the raters were provided with UCOT v.2017-rev1, the draft Guidance Document, and Data Collection Workbook (an entry-validated spreadsheet designed to collect the data; see **Supplemental Materials**). Coders were instructed to use any publicly-available data sources to inform their classification of the record, including but not limited to: the individual’s LinkedIn profile (provided with the data set), employer website, personal website, etc. After the records were coded using UCOT v.2017-rev1, inter-rater reliability analyses were performed, and concordance was found to be insufficient in the Job Function tier (see **Results**).

Following Round 1, the group of six coders met twice by phone to review the data, discuss sources of discordance (see **Methods *Analysis of Discordance***), and develop refinements to the Guidance Document and the taxonomic categories and thereby improve consistency in interpretation. These meetings resulted in UCOT v.2017-rev2, which was tested in Round 2 of the experiment. A new set of alumni records were selected and coded by a group that included 5 experienced coders (from Round 1) and 3 naïve coders who were new to the project. New coders were intentionally recruited to participate in the experiment to explore whether the refinements were sufficient to increase the reliability of interpretation of the documents by an inexperienced user. Inter-rater reliability analyses were performed on Round 2 data, and concordance was found to meet our minimum acceptable Krippendorff’s alpha metric across all three tiers of the taxonomy.

Following Round 2, the group of eight coders met twice by phone to discuss the specific sources of low concordance in UCOT v.2017-rev2. Refinements were made to the taxonomy and Guidance Document, generating the finalized UCOT v.2018 and Guidance Document v.2018.

All raters in both rounds of the experiment were university administrators in professional roles focused on career development for PhD scientists, with experience ranging from 6 months - 15 years in the field. Eight of the nine raters involved in the experiment received doctoral degrees in the biological or social sciences; the team consisted of seven women and two men. The team of raters involved in Round 1 included four experienced raters who had participated in the development of the taxonomic categories and were responsible for design of this experiment; Round 1 coding team also included two naïve raters who were new to both the taxonomy and the experiment. The testing of UCOT v.2017-rev2 in Round 2 involved five experienced raters (now including two who had been naïve in Round 1) and three new naïve raters who had had no previous involvement in the project.

### Selection of Alumni Records

The pool from which the data sets were selected included 2,587 graduate student alumni records which had previously been coded according to the UCOT v.2017 by one individual at each originating institution (*Vanderbilt University School of Medicine, Emory University, and UNC Chapel Hill*). Each record was composed of a unique record number, current job title, current employer, LinkedIn profile or other job-related URL, and graduation date. Postdoctoral alumni were not included in the data set due to inconsistent collection of career outcomes for this population across institutions. For a more detailed description of how alumni records were selected for inclusion in Round 1, please refer to **Supplemental Table 4**.

The final Round 1 data set contained 572 records, including a total of 185 records from UNC Chapel Hill, 192 from Vanderbilt, and 195 from Emory. Women comprised 47% of the dataset, men 28%, and 25% of the alumni records were of unknown gender. **Figure 2** shows the distribution of alumni records according to year of graduation.

**Figure 2:**
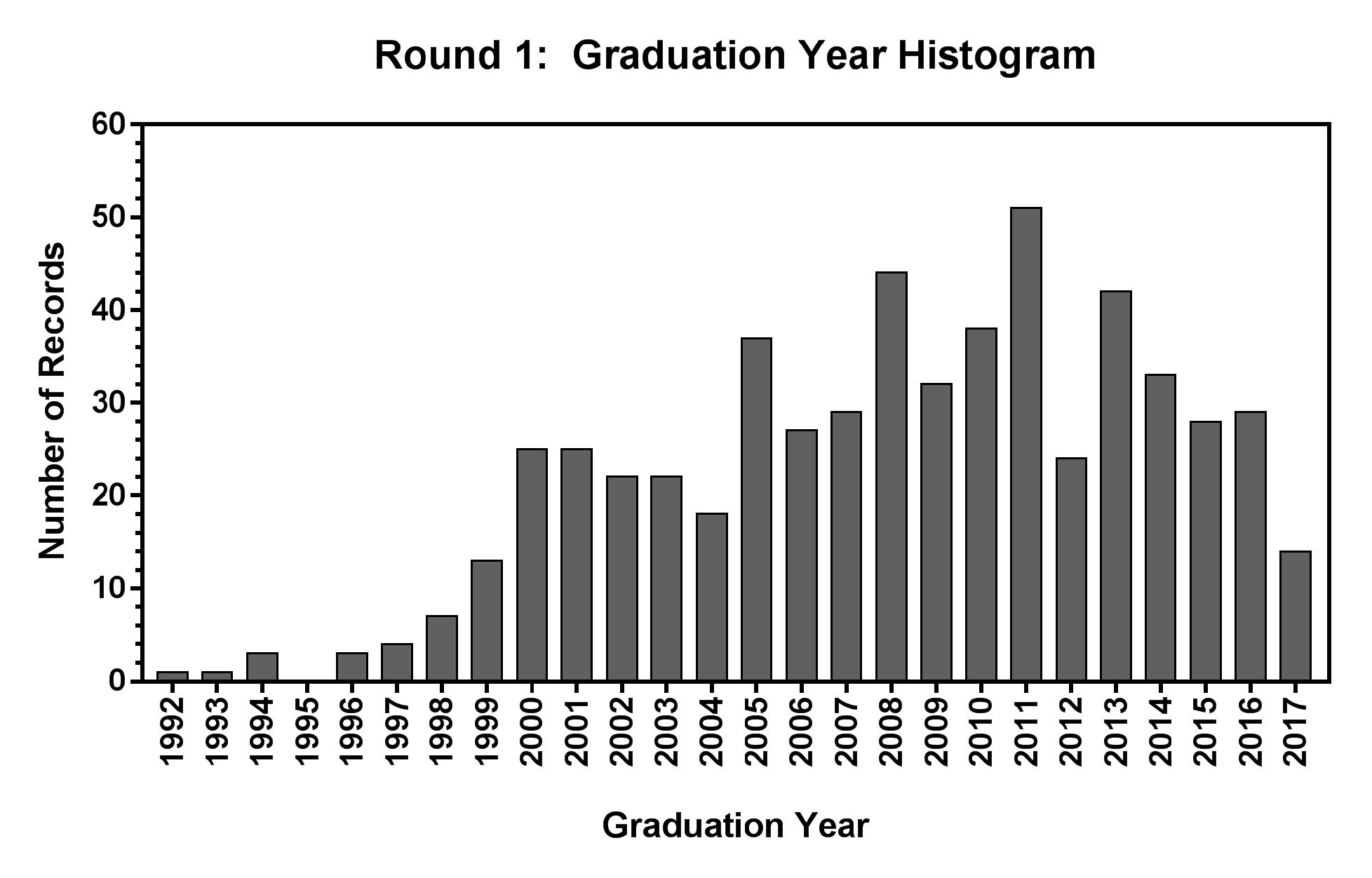
Histogram of Graduation Years for Round 1 Records. Distribution of graduation years for all records coded in Round 1 using UCOT v. 2017-rev1.

Revisions to UCOT v.2017-rev1 were assessed by using this version (UCOT v.2017-rev2) to classify a second data set of 219 records, generated in the same manner as Round 1. In this case, a minimum representative sample across each Job Function category was used to select the number of records; where possible, three records per job function were chosen from each institution. Nine records per job function provide sufficient data to query inter-rater reliability without placing an undue time burden on the Round 2 coders. The Round 2 dataset was comprised of 49% women, 26% men, and 25% unknown. **Figure 3** shows the distribution of records according to year of graduation. For a more detailed description of how alumni records were selected for Round 2 of coding, please refer to **Supplemental Table 5**.

**Figure 3:**
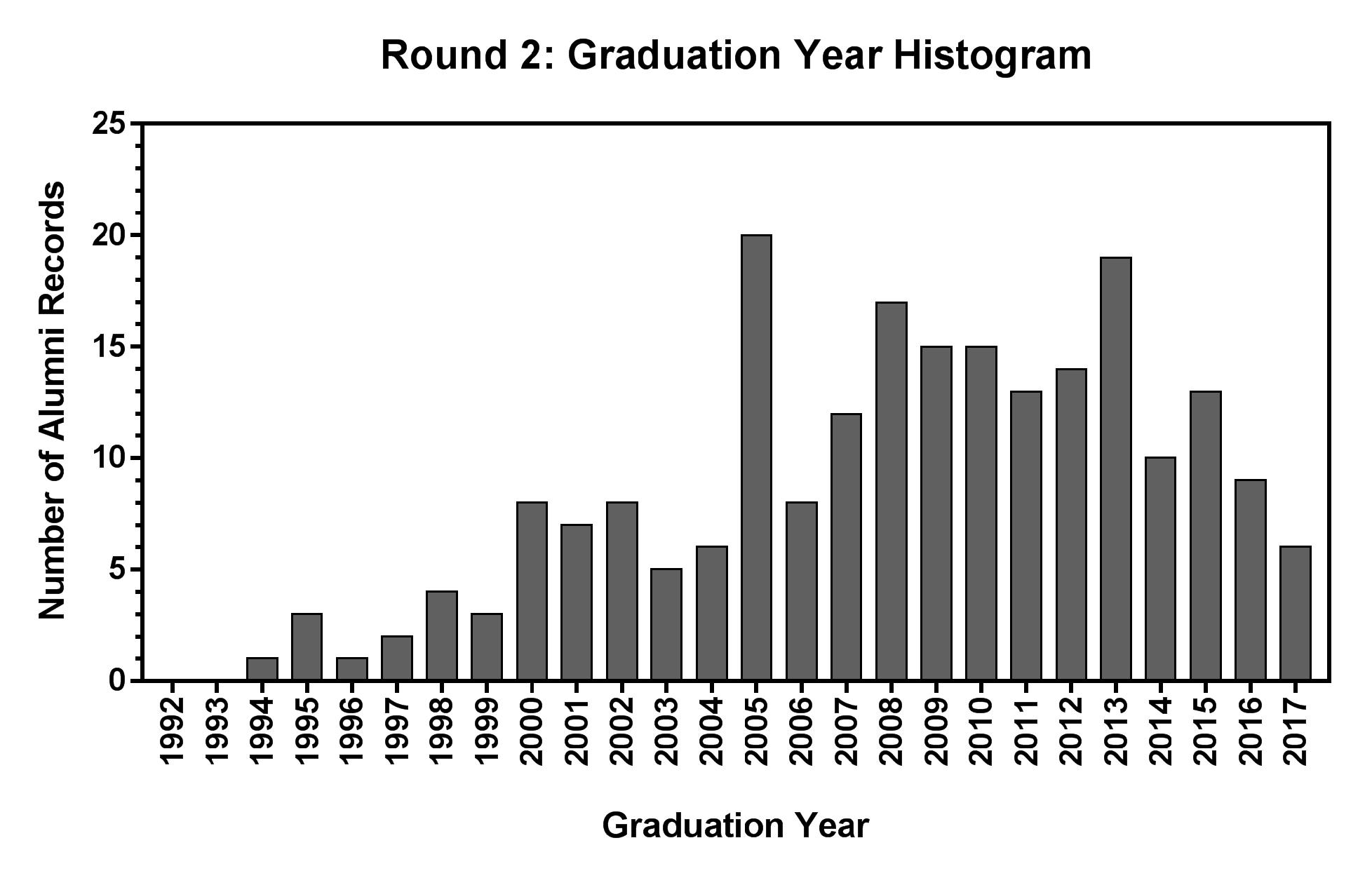
Histogram of Graduation Years for Round 2 Records. Distribution of graduation years for all records coded in Round 2 using UCOT v. 2017-rev2.

### Materials

In Round 1 of the experiment, coders were provided with three documents: 1) a draft Guidance Document for how to code challenging records (*generated by CAS and TH, and piloted at Emory prior to this study*); 2) the taxonomic categories with definitions; and 3) a Data Collection Workbook containing the 572 records to be coded using data-validated fields for classification in each tier. Each record was composed of a unique record number, job title, current employer, LinkedIn profile or other job-related URL, and graduation date. The coding of each record in the workbook was achieved by selecting from drop-down menus of categories in each taxonomic tier (Workforce Sector, Career Type, and Job Function) and a prompt to indicate (yes/no) whether the coder had accessed the provided LinkedIn profile. The draft Guidance Document addressed issues that had been discovered during preliminary use of UCOT v. 2017, including some elaboration on how to classify faculty, entrepreneurs, postdocs and other types of further training positions.

In Round 2 of coding, coders were provided with revised versions of the same three documents: 1) an expanded Guidance Document; 2) the taxonomic categories with definitions where changes had been incorporated; and 3) a Data Collection Workbook containing the 219 records to be coded using data-validated fields for classification in each tier. In Round 2, records in which the employer was an academic institution were augmented with the Carnegie Classification [27] of that institution. Three new open-answer coding fields were added to the Data Collection Workbook, including: 1) drop-down menus related to collecting Faculty Flag data (*if relevant, as discussed below*); 2) prompts to note whether additional sources had been accessed (*e.g., institutional website, personal lab website, etc.*,); 3) prompt to indicate how much time was spent coding the record in minutes.

The Round 2 Guidance Document was significantly expanded and refined from the draft version used in Round 1; modifications included: a) a formal introduction to the taxonomy classification system and recommendations for implementation; b) a set of frequently asked questions (FAQs); c) the taxonomic categories with refined definitions; and d) the list of Carnegie Classifications of Academic Institutions. The guidance FAQs addressed and provided proscriptive clarification for how to code types of jobs that had been identified as problematic in Round 1, as well as explanations for the logic behind the recommendations. Examples included how to:

- Code training positions (*e.g., Fellows, Scholars, Residents, Interns*);
- Distinguish between Principal Investigators and Research Staff;
- Define ‘Entrepreneur’;
- Assign a primary job function for individuals with multiple roles;
- Differentiate For-Profit from Nonprofit entities;
- Differentiate Academic from Nonprofit hospitals;
- Resolve assignment of faculty members to Career Type “Primarily Research” or “Primarily Teaching”, and;
- Implement the Faculty Flag system.

The list of Carnegie Classifications was included for reference, if needed, in the classification of Career Types “Primarily Research” or “Primarily Teaching” (see **Discussion**). We provide summaries of the finalized tiers, categories, and definitions of UCOT v.2018 in **Tables 1, 2, and 3**. Please see **Supplemental Table 1** to view the complete and unabridged UCOT v.2018.

**Table 1:** UCOT v.2018 - Tier 1 Workforce Sector. An individual’s Workforce Sector generally reflects the type of company or institution where they are employed. The unabridged version of UCOT v.2018 is available in **Supplemental Table 1**.

| WORKFORCE<br>SECTOR | Definition | Coding clarifications |
| --- | --- | --- |
| **Academia** | Any post-secondary academic institution where training occurs, including colleges, universities, some medical centers, or freestanding research institutions. | Includes teaching hospitals.<br>Does not include K - 12 schools. |
| **Government** | Any organization operated by federal, state, local or foreign governments. | Includes Veterans Affairs hospitals and some K - 12 schools. |
| **For-Profit** | Any organization that operates to make a profit, including some industry research. |  |
| **Nonprofit** | Any non-governmental organization that does not operate to make a profit. |  |
| **Other** | Individuals who work at an organization not included in other options or who are on extended leave of absence from the workforce. | Includes individuals known to be unemployed. |
| **Unknown** | No current career information is known. | Reserve for individuals without any title or any available employment information. |

**Table 2:** UCOT v.2018 - Tier 2 Career Type. An individual’s Career Type should reflect the general content of their work. The unabridged version of UCOT v.2018 is available in **Supplemental Table 1**.

| **CAREER TYPE** | **Definition** | **Coding clarifications and examples** |
| --- | --- | --- |
| **Primarily Research** | The primary, although not necessarily only, focus is the conduct or oversight of scientific research. | See guidance document for suggestions on how to code faculty titles. *Includes postdoctoral research positions. |
| **Primarily Teaching** | The primary, although not necessarily only, focus is education and teaching. | See guidance document for suggestions on how to code faculty titles. *Includes postdoctoral teaching positions. |
| **Science-Related** | Career that is relevant to the conduct of scientific research, but does not *directly* conduct or oversee research activities. | Program Officer; Physician; Medical Science Liaison; Editor; Healthcare Consultant |
| **Not Related to Science** | Career that is not directly relevant to the conduct of scientific research. | Bank Manager; Campaign Manager; Golf Instructor; Chef; Painter |
| **Further Training or Education** | Temporary training position or enrollment in further education. | Law School Student; MD/PhD Student; AAAS Fellow |
| **Unemployed/Unknown** | Not currently employed or no information is known. | Family caretaker; retired; unemployed. |

**Table 3:** Unified Career Outcomes Taxonomy v.2018 - Tier 3 Job Function. An individual’s Job Function is defined by specific skill sets and/or credentials required for employment in that function. The table as represented here is significantly abbreviated and omits two columns that are new to UCOT v.2018: “Job Title Examples” and “Standard Coding Schema.” The unabridged version of UCOT v.2018 is available in **Supplemental Table 1**.

| **JOB FUNCTION** | **Definition** | **Coding clarifications** |
| --- | --- | --- |
| **Administration** | Administrative-intensive roles in which the individual is managing people, projects, and resources. | Spends more than 50% of their time in administrative duties that are internal to the organization. |
| **Business Development, Consulting, and Strategic Alliances** | Role relevant to the development, execution, management, or analysis of a business. Role may include external relationship management, refinement of operational efficiency, or fee-based advisory services. | Director or higher-level positions that have primarily *externally-facing* responsibilities |
| **Clinical Research Management** | Role that is responsible for the oversight, management, or design of clinical research trials. |  |
| **Clinical Services** | Role that involves that administration of clinical services or research. |  |
| **Data Science, Analytics, and Software Engineering** | Role that primarily involves programming, analytics, advanced statistics, data communication, and/or software development. |  |
| **Entrepreneurship** | Founder or co-founder of their own business or enterprise, and employer of at least one other person. | This category should not be used for C-suite executives of established companies, they should be classified as "Business Development" unless they were also involved in founding the company. Freelance specialists should be classified in the field of their specialization (*e.g., science communications, consulting*). |
| **Group Leader or Principal Investigator** | Role that involves leading a research team in any research environment. The individual is clearly responsible for funding and managing a research group. | This job function will be used in multiple sectors (academia, nonprofit, for-profit, government). |
| **Healthcare Provider** | Role where the primary responsibility is providing healthcare. | These individuals usually fall into the Career Type "Science-related," unless it is clear that they are primarily performing research; in that case, the career type might be "Primarily Research." |
| **Intellectual Property and Law** | Role that involves the curation, management, implementation or protection of intelligence and creation, including patents, trademarks, copyrights, or trade secrets. | Law students belong to this Job Function (instead of "Completing further education") because their career outcome is specific. Their training status will be captured at the Career Type level as "Further Training." |
| **Postdoctoral** | A temporary mentored training position following completion of doctoral degree. | At the Career Type level, classify the postdoc according to the type of postdoctoral research that they are engaged in, likely either "Primarily Research" or "Primarily Teaching." |
| **Regulatory Affairs** | Role that involves controlling or evaluating the safety and efficacy of products in areas including pharmaceuticals, medicines, and devices. |  |
| **Research Staff or Technical Director** | Role that directly involves performing and managing research. | The individual is involved in research, but not responsible for funding and managing a group. "Staff fellow" is coded here if it is not an official postdoctoral position. |
| **Sales and Marketing** | Role that is related to the sales or marketing of a science-related product or service. | A medical science liaison should be classified as "Technical Support and Product Development." |
| **Science Education and Outreach** | Role that involves K-12 teaching or public outreach at a primary/secondary school, science museum, scientific society, or similar. | A public school belongs in the Government Sector. |
| **Science Policy and Government Affairs** | Role that involves policy or program development and review, including analysis, advisory, or advocacy. | Unless a program officer is actively involved in determination of policy, that individual should be classified as "Administration." |
| **Science Writing and Communication** | Role that involves the communication of science-related topics. |  |
| **Teaching Faculty or Staff** | Teaching position at post-secondary schools, universities, or institutions with no contractual research responsibilities. | Positions in elementary and high schools should be coded as "Science Education and Outreach." |
| **Technical Support and Product Development** | Role that requires specialized technical/expert knowledge of a science-related product. |  |
| **Other Employment** | Role that does not require scientific training or involve the direct implementation or communication of science. | Includes: Industry not related to science; politics not focused on science; food or hospitality services; military service; volunteer/mission work. |
| **Completing Further Education** | Pursuing additional education that usually results in graduation with conferment of a degree or certificate; this does not include traditional academic postdoctoral positions which have their own Job Function "Postdoctoral." | Pursuing additional education where the career outcome is not specific or clear. Postdoctoral trainees should be classified in the "Postdoctoral" Job Function and Career Type based on their broad field. |
| **Unemployed or Seeking Employment** | Known to be temporarily out of the workforce but likely to return. |  |
| **Deceased/Retired** | Permanently out of the workforce. |  |
| **Unknown** | No title or career information can be found for this individual. |  |

### Analysis of Discordance

We employed two methods to identify discordance following Round 1 of coding. Both methods share a common starting point, which was to determine the Workforce Sector, Career Type, and Job Function that was most commonly chosen (the mode) by the coders for every record. The mode was then employed as the presumed “correct” answer. To calculate the average number of unique categories per tier for each record (*columns A and C in* **Supplemental Table 6**), we counted the number of unique categories that coders applied to the given record. We then calculated the average number of unique categories across all records in each tier (as defined by the most popular choice). Percent discordance was measured by determining, for every record, how many coders did *not* choose the same category as the mode. Subsequently, that number was divided by the total number of coders and multiplied by 100 to give a percentage (*columns B and D of* **Supplemental Table 6**).

## RESULTS

Based in preliminary data not included here, we initiated the experiment by modifying the UCOT v.2017 (*available for download* [RBR website]) to clarify known sources of error, therein generating UCOT v.2017-rev1. This version was used to code 572 records in Round 1 of the experiment. Using a Krippendorff’s Alpha of 0.70 as the minimal threshold to define reliability among six coders (see **Methods *Procedure***; [23, 24]), we observed that the Workforce Sector tier (0.83 agreement) and the Career Type tier (0.70 agreement) met the acceptable parameters of reliability (**Figure 4**). However, the measured reliability for the Job Functions tier (0.62 agreement) was below admissible levels (**Figure 4**).

**Figure 4:**
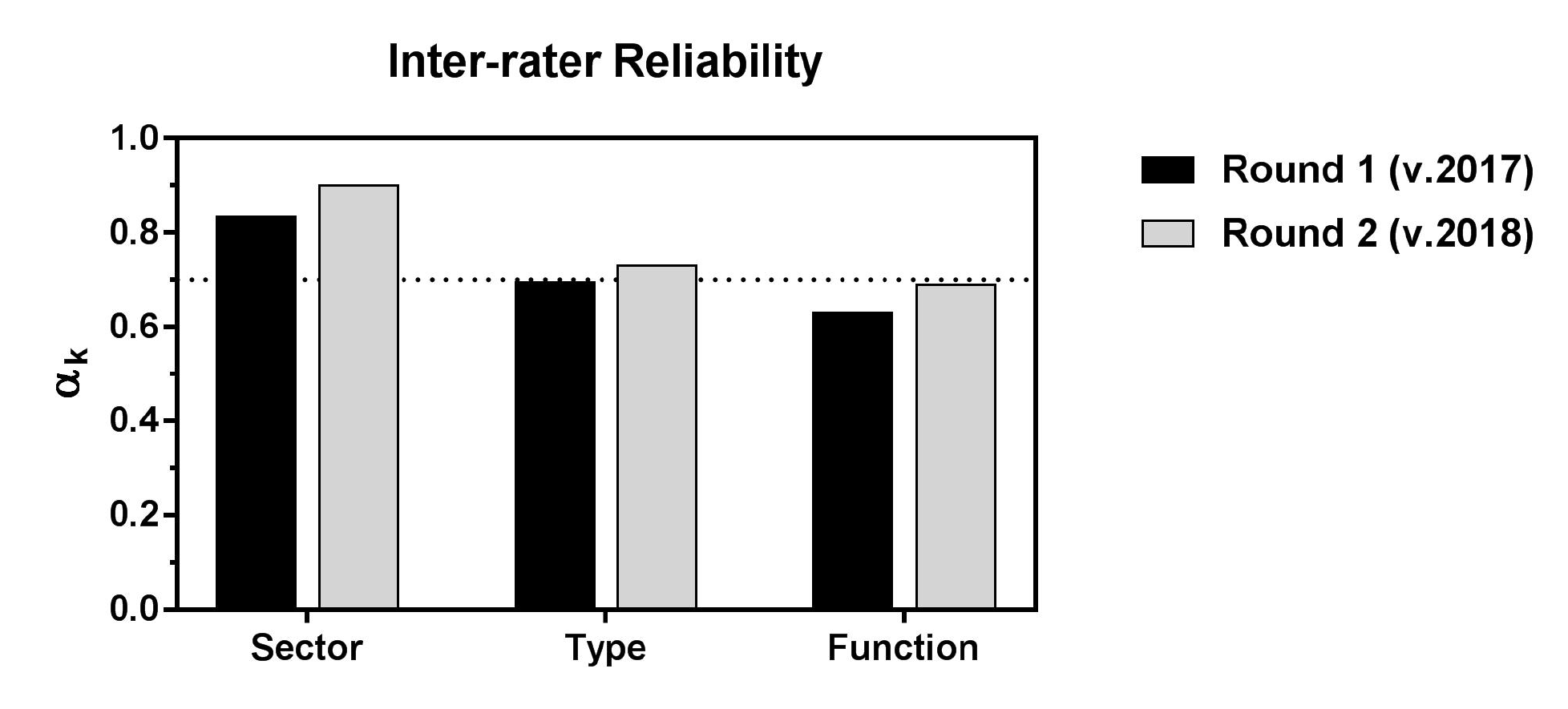
Inter-rater reliability scores for Round 1 (UCOT v.2017-rev1) and Round 2 (UCOT v.2017-rev2) of coding, as measured by Krippendorff’s alpha (α_k_). Workforce Sector, Career Type, and Job Function were coded by six coders in Round 1 (black bars) and eight coders in Round 2 (gray bars). The threshold is indicated by the horizontal dotted line. Round 2 results for every category are higher than Round 1. Coders used 6 categories for Workforce Sector, 6 categories for Career Type, and 26 (Round 1) or 23 (Round 2) categories for Job Function.

The six coders in Round 1 had six-way agreement on 77% of all records within Workforce Sector, 55% of all records within Career Type, and 36% of all records within Job Function. Furthermore, we reached five-way agreement on 95% of records for Workforce Sector, 95% of records for Career Type, and 73% of records for Job Function, suggesting that, in many cases, only one person interpreted a definition differently than the rest of the group.

To identify hot spots of discordance, we ranked the records by number of unique categories utilized by the coders and looked for patterns among records that had high disagreement (an illustration of the discordance is provided in **Supplemental Table 6**). Within the Career Type tier, we identified discordance in application of the “Further Training” category and in differentiating “Primarily Research” and “Primarily Teaching” faculty positions. Job Functions with high levels of discordance included: “Faculty-Non-Tenure track”; “Clinical Services”; “Sales and Marketing”; “Group Leader”; “Business Development, Consulting, and Strategic Alliances”; “Clinical Research Management”; and “Administration”.

While we hypothesized that most discordance issues could be addressed by providing additional examples in the Guidance Document or by making minor clarifications in taxonomic definitions, we also hypothesized that two structural changes to UCOT v.2017-rev1 would improve reliability. To address the discordance in the coding of the five faculty Job Functions in UCOT v.2017-rev1 (“Adjunct/part-time teaching staff”, “Faculty: non-tenure track”, “Faculty: tenured/tenure track”, “Faculty: tenure track unclear or not applicable”, and “Instructor/full-time teaching staff”), the group agreed to test an alternative approach to capturing faculty data, originally developed by the Biomedical Research Education and Training office (BRET) at Vanderbilt (*KP and AB, unpublished*). Specifically, in addition to classifying each record by the three primary taxonomic tiers, the record is also examined for any indication in the job title that the individual holds a faculty designation at their employing institution. If so, the record is ‘flagged’ as faculty and additional information is collected, including title (*e.g., professor, instructor, lecturer*), rank (*e.g., assistant, associate*), and prefix (*e.g., adjunct, research, teaching*). This strategy for capturing faculty status permits us to combine the five faculty-related Job Functions in UCOT v.2017 and UCOT v.2017-rev1 into just two Job Functions in UCOT v.2017-rev2: “Principal Investigator or Group Leader” and “Teaching Faculty or Staff.” Examples of the Faculty Flag schema with sample data are provided in the Guidance Document (**Supplemental File 3**). The second structural change was introduced in the Career Type tier by adding an “Other” category to more accurately capture alumni for whom some information is known, to avoid classifying them as “Unknown”. Incorporation of these changes resulted in the UCOT v.2017-rev2, which was then used in Round 2 by a second group of coders to code a new dataset of 219 records (see **Methods**). Notably, the resultant Krippendorff’s alpha values demonstrated improved inter-rater reliability in this second experiment (**Figure 4**).

We considered that the elevated reliability achieved in Round 2 using UCOT v.2017-rev2 may have resulted from the coders’ greater familiarity with the taxonomy, developed through Round 1 of coding and subsequent conversations about sources of discordance. To explore this possibility, Round 2 reliability was calculated separately for experienced and naïve coders. Although experienced coders performed slightly better, naïve coders did not lag far behind (**Figure 5**).

**Figure 5:**
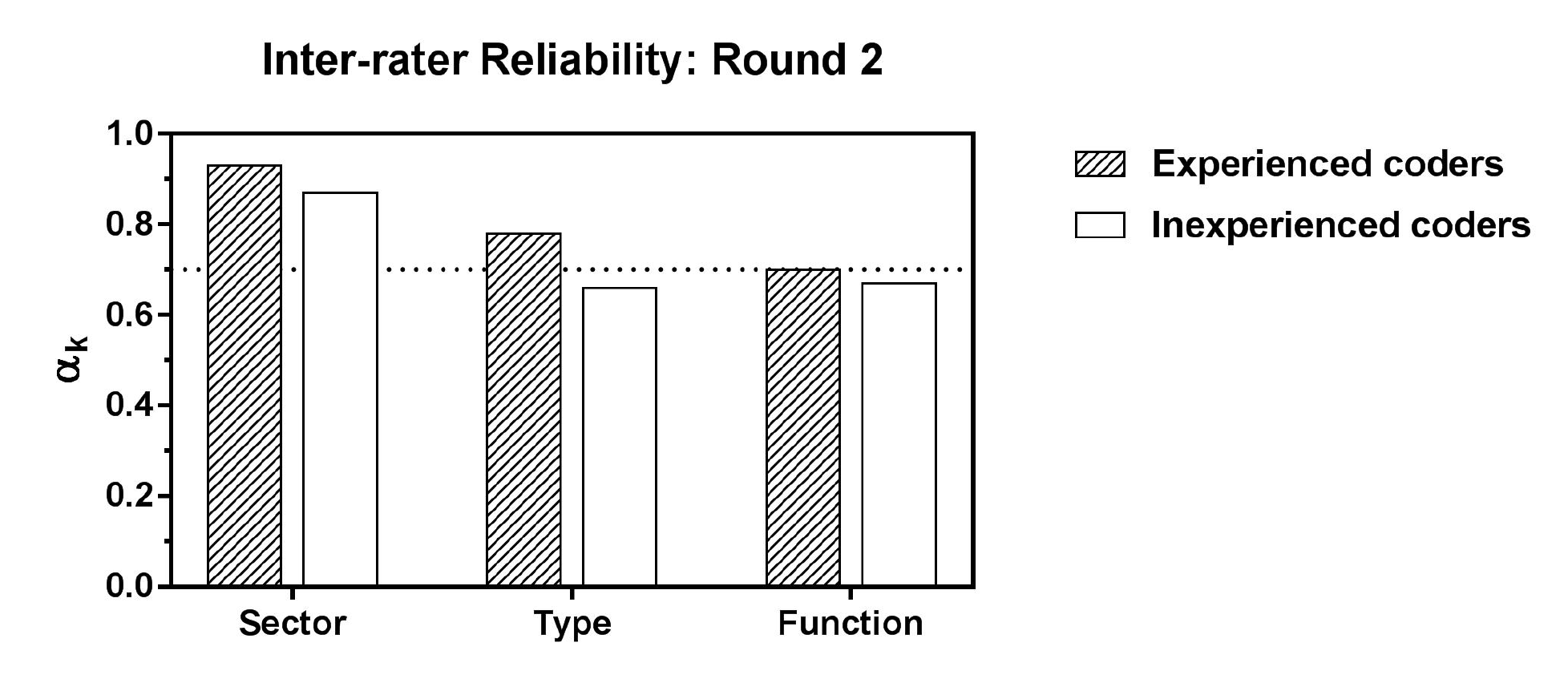
Inter-rater reliability scores (αk) for experienced and naïve coders in Round 2 of coding. Workforce Sector, Career Type, and Job Function were coded by and eight coders in Round 2. The results are similar for experienced (n=5, striped bars) and naïve coders (n=3, white bars). Coders used 6 categories for Workforce Sector, 6 categories for Career Type, and 23 categories for Job Function.

The introduction of naïve coders to the Round 2 experiment also permitted us to explore whether the amount of time spent coding each record significantly differed between naïve and experienced coders. Therefore, in Round 2, all coders were prompted to record how much time was spent in coding each record. The total time for experienced coders ranged from 213 to 540 minutes (average 328 minutes) and 246 to 545 minutes for naïve coders (average 429 minutes). The overall average time to classify the second round of 219 records was 366 minutes, which equates to about 1 minute, 40 seconds per record.

## DISCUSSION

In this study, we tested the consistency of categorizing the records of PhD alumni using a collaboratively-established, three-tier career outcomes taxonomy. Using Krippendorff’s alpha to measure the inter-rater reliability, we found that consistency among coders was initially achieved for the first two taxonomic tiers, but not for the Job Function tier. To improve inter-rater reliability, we identified several sources of discordance, and using these results, tested a revised taxonomy that included clarifying definitions and a guidance rubric. After iterative rounds of coding, we achieved acceptable levels of reliability among all three tiers.

We ascribe the majority of the improvement in reliability during Round 2 (**Figure 4**) to the provision of a detailed and proscriptive Guidance Document (**Supplemental File 3**). The value of this document lies in its development through classification of 2500+ alumni records and the collective reflection of experts in the field on the nuances of these records. We intentionally enriched the data set with less common Job Functions to ensure full coverage of the entire taxonomy, and thereby uncover potentially uncommon types of employment that were poorly defined or described in the taxonomy itself. While there are likely to be unique forms of employment that we did not encounter in our datasets and therefore are not addressed in the Guidance Document v.2018, we are confident that we obtained full coverage of the most common career outcomes for PhDs in the biosciences and that the Guidance Document v.2018 provides sufficient instruction to reliably code these career outcomes.

Our data indicate that institutions may consider the value of training and retaining experienced coders year after year; however, the use of inexperienced coders should not preclude an institution from starting to classify alumni career outcomes (**Figure 5**). As coders are first trained, we recommend that institutions consider internally measuring and evaluating inter-rater reliability on sample records, *regardless of which taxonomy they choose to employ*. To that end, institutions or academic divisions may find that it is most efficient to designate a small, centralized team to code all alumni records, as opposed to approaching classification in a more decentralized manner that involves many naïve coders.

The results from this study reveal that clarifying modifications to definitions can have a significant effect on the consistency of interpreting taxonomic categories, which ultimately will affect how jobs are classified by individual coders. Therefore, we include a summary of the clarifying modifications we made to UCOT v.2017, in the hopes that the proposed updated definitions and categories of UCOT v.2018 will enhance cross-institutional reliability and reproducibility of data classification. An annotated table of all changes made to UCOT v.2017 in generating UCOT v.2018 is available in **Supplemental Table 2**.

## SUGGESTIONS TO IMPROVE THE RELIABILITY OF THE UNIFIED CAREER OUTCOMES TAXONOMY v.2017

### Tier 1: Workforce Sector

In the Workforce Sector tier, we suggest that Academia be defined as “any post-secondary academic institution where training occurs, including colleges, universities, some medical centers, or freestanding research institutions.” We added the specification that K-12 teaching jobs be classified as belonging to Government, Nonprofit, or For-Profit sectors, instead of Academia.

### Tier 2: Career Type

In the Career Type tier, we suggest adding the term ‘unemployed’ to the existing category “Unknown,” renaming this category “Unemployed/Unknown.” In the absence of this category, individuals who are not currently employed— but who are known to be searching for a job or intend to return to the workforce— could only be classified as Career Type “Unknown,” thereby losing information that may actually exist about their current employment status. Now, by classifying them as “Unemployed/Unknown” at the Career Type tier and combining that with a new Job Function category, “Unemployed or Seeking Employment” (*described below*), we are able to capture current status of individuals who have self-reported that they are, for example, acting as family caregivers or on the job market.

Second, our group recognized the need to develop guidance for how to classify academic faculty members in the Career Type tier, where the likely options are “Primarily Research” and “Primarily Teaching.” We acknowledge that this classification is complicated by the multi-functional roles played by many faculty members and that a large proportion of academic faculty balance both research and teaching. We recommend that the coder use all available public resources to determine in which capacity the faculty member is likely to spend most of their time. Examples of this type of data include but are not limited to: frequency and recency of peer-reviewed research articles (e.g., publication record on PubMed); information related to the number of courses that an individual taught in a given academic year; and, supervising the research of doctoral students or postdoctoral scholars. If no illuminating data can be found, we recommend that a coder use the Carnegie Classification [27] of the employing institution to determine Career Type: if the individual is a faculty member at an academic institution with a Carnegie Classification of 15 or 25 (“highest research activity”), categorize the individual as “Primarily Research”; faculty at institutions in any other Carnegie Classification should be categorized as “Primarily Teaching.” (See further examples of this coding strategy in the UCOT v.2018 Guidance Document, which is **Supplemental File 3**.) If at all possible, we recommend *against* using the Carnegie Classification strategy due to its potential inaccuracy. This default coding will misrepresent some faculty as “Primarily Research” because many institutions with Carnegie Classification 15 or 25 are known to appoint faculty members to serve in primarily teaching roles who bear little to no research responsibilities. This default strategy will also underreport the research responsibilities held by many faculty members at institutions with Carnegie Classification other than 15 or 25. A related challenge is classifying the tenure status of a faculty member; we address this in detail in our recommendations for modifications to Job Functions below. Because of these challenges, we recommend implementation of a Faculty Flag (discussed in detail below) and propose further elaboration of the Faculty Flag strategy as a potential solution for future investigation.

Third, we propose a change in the way that postdoctoral scientists are classified in the Career Type tier. We suggest that they be classified as “Primarily Research” or “Primarily Teaching” instead of as “Further Training or Education” (*as defined in UCOT v.2017*), since their postdoctoral training status is already captured in the Job Function tier. We make this suggestion due to our community’s desire to understand the true number of research-focused postdoctoral scientists involved in the national workforce [6, 28, 29]. Currently in UCOT v.2017, postdoctoral scientists are classified as “Further Training or Education” at the Career Type tier, followed by “Postdoctoral” at the Job Function tier, therein losing the specific nature of their postdoctoral work. For example, if the individual is engaged in a postdoctoral training program with a significant teaching focus, the UCOT v.2017 taxonomy cannot capture the teaching component in their training. Following our suggestion in UCOT v.2018, that individual would be classified as “Primarily Teaching” at the Career Type tier and as “Postdoctoral” at the Job Function tier and we will have more detailed understanding of the *nature* of their postdoctoral experience.

As a corollary, we suggest that the “Further Training or Education” category within the Career Type tier be reserved explicitly for specialized training experiences *distinct from PhD education*; examples include postdoctoral fellowships in governmental policy (*e.g., American Association for the Advancement of Science (AAAS), Science and Technology Policy Fellowship*) or academic programs that will lead to an additional degree or certification (*e.g., law school, MD/PhD students in second stage of training*). We suggest that the focus of this further training can be captured at the Job Function tier. For example, a student in law school would be classified as “Academia → Further Training or Education → Intellectual Property and Law.” Similarly, a AAAS Fellow would be classified as “Nonprofit → Further Training or Education → Science Policy and Government.” Other unique fellowships, such as through the ORISE programs, will require careful attention during coding to accurately reflect the content of their work. Additional examples to help coders determine how to apply Job Function categories to postgraduate and postdoctoral training programs are offered in the Guidance Document (**Supplemental File 3)**.

We recognize that institutions that are currently classifying postdoctoral scientists as Career Type “Further Training or Education” are doing so to highlight the temporary training nature of the postdoctoral position. We also recognize the necessity of classifying postdocs as “Further Training or Education” if an institution is only collecting data for tiers 1 and 2 (Workforce Sector and Career Type); institutions otherwise might grossly undercount the number of postdoctoral scholars. However, for those institutions that are currently or plan to collect data for the 3rd tier (Job Function), our concern is that by classifying postdocs using the “Further Training or Education” followed by “Postdoctoral” categories, we *lose resolution about the very population that we hope to track and analyze more precisely.* As a consequence, we also lose valuable opportunities to discover distinct shifts among PhD outcomes, such as identifying longitudinal trends in the types of postdoctoral training experiences pursued by our trainees (*e.g., a decrease in number of research-focused postdocs accompanied by a rise in teaching-focused or government policy postdocs etc.*).

Though we propose this as one approach for using UCOT to provide detailed information about the nature of postdoctoral training experiences, we acknowledge that there are other adaptations of UCOT that could be considered. Alternative strategies might include: (1) creating additional Job Functions to capture different types of postdoctoral positions (*e.g., “Postdoctoral: Research” and “Postdoctoral: Teaching”*); (2) pulling all postdoctoral and training categories out of Career Type and Job Function, classifying trainees in Career Type and Job Function categories most similar to their type of work and adding a “Training Flag” (denoting “Student” or “Postdoctoral”); (3) creating a fourth taxonomic tier focused on level of seniority or autonomy in a role (*e.g., Student, Postdoctoral, Staff, Director*); or (4) creating a “Postdoc Flag” to capture the focus of their postdoctoral position (e.g., research, teaching, policy, etc). In the latter case, the individual would be classified as Career Type “Further Education or Training” (as is the current recommendation in UCOT v.2017), followed by Job Function “Postdoctoral,” and an additional data collection field would prompt the coder to select the focus of the postdoctoral position (*research, teaching, policy, etc.).* We successfully employed a “flag” approach in the classification of faculty so that we could simultaneously capture the diversity of faculty responsibilities and increase the reliability of categorization among raters; this is discussed in more detail below. Regardless of the approach an institution chooses to employ, we encourage institutions to consider how to most accurately represent the composition of the postdoctoral populations within their dataset.

### Tier 3: Job Function

We identified three important clarifications to definitions in the Job Function tier, and one larger structural modification; implementing these changes in UCOT v.2017-rev2 improved concordance among raters within the Job Function tier. Below we summarize these modifications; all updated definitions and notated changes are provided in **Supplemental Table 2**.

First, we suggest the addition of a new Job Function category “Unemployed or Seeking Employment” to more accurately classify individuals for whom some information is known about their employment status (*e.g., individuals who are known to be family care providers or seeking employment).* Currently in UCOT v.2017, these individuals would be classified as Job Function “Unknown.”

Second, we found that our group faced significant challenge in distinguishing between the Job Functions “Administration” and “Business Development, Consulting, and Strategic Alliances”. Upon further discussion, we concluded that discordance resulted from specific job titles such as “Chief Financial Officer” or “Academic Dean” that relate to the administration of a company/institution (and therefore might be classified as “Administration”) yet entail significantly more responsibility and a different skill set than a departmental administrator or program manager. We suggest that these Job Functions could be differentiated by the *primary focus of their work* (defined as 50% or more of an individual’s time spent on these duties): strictly administrative roles spend most of their time in duties that are *internal to the organization*, while business development roles are generally higher-level positions that have *externally-facing responsibilities*. Under this revised definition, a dean of the biological sciences division would be classified as “Academia → Science-Related → Business Development, Consulting, and Strategic Alliances,” while a departmental administrator would likely be classified as “Academia → Science-Related → Administration.” Extrapolating to other Workforce Sectors, a project review manager at a For-Profit industry company would be classified as “For-Profit → Science-Related → Administration,” while the director of medical affairs at the same for-profit industry research company would be classified as “For-Profit → Science-Related → Business Development, Consulting, and Strategic Alliances.” In the Government Sector, a grant administrator at National Institutes of Health would likely be classified as “Government → Science-Related → Administration,” and the director of the National Institutes of Health would be classified as “Government → Science-Related → Business Development, Consulting, and Strategic Alliances.” We recognize that the individual tasked with classifying alumni records may not be able to distinguish the full spectrum of duties involved in a specific job and may feel ill-equipped to categorize these individuals. We encourage coders to use public data sources to their fullest extent, accepting that, as in all projects that are based in subjective assessment, there will be some degree of error introduced by interpretation.

Third, we identified discordance generated by job titles that included ‘medical science liaison’, which had been defined in UCOT v.2017 as belonging in the “Sales and Marketing” Job Function. We quickly recognized that, even within our group of career development professionals, there were different understandings of what a medical science liaison actually does on a day-to-day basis; this is an example of a job that has evolved significantly over the past decade and can vary greatly depending on the organization or employer. After significant discussion and outreach to individuals who hold the ‘medical science liaison’ job title, we concluded that, by default, these individuals should be classified in the “Technical Support and Product Development” Job Function. We recognize that there may be exceptions to this rule and encourage coders to obtain as much publicly-available information as possible about the individual’s actual duties to make the most informed categorization for each unique individual.

Our final recommendation in the Job Function tier relates to the classification of academic faculty and teaching staff. In UCOT v.2017, academic faculty are classified according to their tenure status: “Faculty: Tenure track”, “Faculty: Non-tenure track”, or “Faculty: Track unclear or not applicable.” During preliminary use of UCOT v.2017 (not reported here), multiple classification strategies emerged among the raters. Some coders assumed tenure status based on the institution’s Carnegie Classification (R1, R2, etc.) and lack of an ‘adjunct’ preface to their faculty title, resulting in the coding of nearly all faculty positions as “Faculty: Tenure track.” Other coders concluded that a tenure-track assignment could not be made without direct confirmation from the individual, given that many institutions appoint non-tenure track assistant professors, or have eliminated the traditional tenure structure altogether [19]. This non-standardized interpretation of the categories resulted in significant discordance in the classification of faculty records. Therefore, in the draft Guidance Document that accompanied UCOT v.2017-rev1 during Round 1 of the experiment (*see* **Figure 1** *for experimental procedure*), coders were given explicit instruction to *not* code faculty into the “Faculty: Tenure-track” category unless they were absolutely certain that the individual held a traditional tenure track faculty appointment. This resulted in no application of that category during Round 1 (**Supplemental Table 6**). Even so, during that round we *still* observed high discordance in application of the categories “Faculty: Non-tenure track” and “Faculty: Track unclear or not applicable” and the determination of Career Type as “Primarily Research” or “Primarily Teaching.”

In light of the inability of multiple career development professionals to consistently and accurately assess the tenure status of an individual based on their job title and home institution, as well as changing attitudes about academic tenure [30], we recommend that individuals with academic titles— such as professor, instructor, adjunct professor, research associate professor— be classified under “Principal Investigator or Group Leader”, “Research Staff or Technical Director”, or “Teaching Faculty or Staff” based on information sourced from an individual’s public web presence (*PubMed, LinkedIn, institutional or personal web pages*). In order to permit finer resolution of the faculty population, we recommend that each faculty record be identified by a Faculty Flag and further notated with their faculty title, rank, and prefix (see **Methods**). In our experiment, adding the Faculty Flag to UCOT v.2017-rev2 greatly reduced discordance in the classification of faculty positions in Round 2 (**Supplemental Table 6**).

Multiple benefits emerged with the implementation of the Faculty Flag in UCOT 2017-rev.2 (for additional examples of the versatility of the Faculty Flag approach, please see **Supplemental File 3**). Five previous faculty Job Functions (“Adjunct/part-time teaching staff”, “Faculty: non-tenure track”, “Faculty: tenured/tenure track”, “Faculty: tenure track unclear or not applicable”, and “Instructor/full-time teaching staff”) were tailored down to two (“Principal Investigator” and “Teaching faculty or Staff”), resulting in a more simplified taxonomy. Furthermore, our raters captured and categorized *more information* from alumni records than just a presumed tenure status. Specifically, this strategy allowed coders to use the Job Function category to describe the functional duties that comprise the majority of a faculty member’s time, without undercounting or misreporting their faculty appointment. For example, UCOT v.2017-rev2 and UCOT v.2018 permit faculty with multiple academic titles, such as deans and directors, to be classified as having a “Business Development, Consulting, and Strategic Alliances” or “Administrative” Job Function, respectively, and simultaneously flagged as faculty. Finally, our decision to remove the focus on tenure from the Job Function categories allowed us to more directly compare functionally similar roles across Workforce Sectors. Specifically, the definition “a role that involves leading a research team in any research environment where the individual is clearly responsible for funding and managing a research group” equally describes a principal investigator at an academic institution and a group leader at an industry research company. Therefore, we suggest that the two be combined into a single category: “Group Leader or Principal Investigator.” The Workforce Sector classification, based on the individual’s employer, will clarify whether the individual performs this function in an academic, for-profit, nonprofit, or government environment. By recognizing the homology that exists between principal investigators in Academia and group leaders in other sectors, UCOT v.2018 underscores the suitability, relevance, and contributions of PhD-trained scientists across the workforce.

We recognize that the collection and classification of records at the granular resolution of Job Function may seem burdensome or unnecessary to some institutions. We also have learned that, while the Job Function data is invaluable to developing career support services internal to the institution, it may be less relevant to national reporting standards. Indeed, the NGLS institutions have agreed to publish the Workforce Sector and Career Type tiers and are not requiring public reporting of the Job Function tier (although most schools are, in fact, collecting that data). However, we strongly recommend the collection of Job Function data, even if it is not publicly disseminated. Job Function data can have great value at both the local, intra-institutional level, and national level. These data provide valuable insights into the types of skill sets used by PhDs. More than Work Sector or Career Type, Job Function can inform the ongoing development of graduate and postdoctoral training support services to address changing needs in the scientific workforce. Job Functions also provide richer data to assist trainees or prospective trainees in making informed career decisions, and the alumni who are known to have pursued specific careers become invaluable resources to current trainees. PhD training develops a versatile skill set, as demonstrated by PhDs who become serial entrepreneurs, government policy advisors, science educators, and public communicators, contributing to society in ways that can be claimed and espoused by the institutions that prepared them to be successful in those ways. Collecting and publicly disseminating the breadth of career paths that PhDs pursue is essential for institutions, policy makers, and the public to assess the impact of their investment in the PhD training enterprise.

## CONSIDERATIONS IN IMPLEMENTING THE UNIFIED CAREER OUTCOMES TAXONOMY

Ideally, adoption of a single taxonomy across all institutions would enable inter-institutional analyses of alumni career outcomes. However, establishing a static and universally-acceptable career outcomes taxonomy across institutions will be challenging. Our results underscore that clarifying modifications to the taxonomy will be needed as more cases of utilization emerge and as the workforce evolves. Some institutions or disciplines may find that some categories are not fully applicable; others may choose to alter definitions or create additional job functions to better suit their needs. For any changes made to the taxonomy, we urge institutions to clearly and prominently annotate these definitional differences when displaying the data, so that others comparing data across institutions will not make false comparisons. Furthermore, we encourage the community to come together periodically to discuss challenging cases, consider recommended changes, and test and refine inter-rater reliability for new definitions. This will be necessary if we are to sustain a career outcomes taxonomy that enables merged and comparative analyses across institutions.

One perceived limitation of the current taxonomy is its specificity to the biological sciences. Interestingly, early tests of this suggest that this taxonomy can be easily adapted for other disciplines. For example, cross-disciplinary implementation of the UCOT v.2017 at Wayne State University identified that, after replacing any occurrence of the word ‘science’ in the taxonomy with the word ‘discipline,’ the existing Career Types and Job Functions were sufficient to classify career outcomes across disciplines (*Ambika Mathur, personal communication*). In practice, in the Career Type tier, changing the category “Science-Related” to “Discipline-Related” or, in Job Functions, changing the category “Science Communications” to “Discipline-Related Communications” made the taxonomy sufficient for cross-disciplinary implementation, with the single exception of adding the category “Author” to the Job Function tier. With more than a dozen institutions already adopting this taxonomy, we encourage continued collaboration across disciplines to further explore its versatility.

A major strength of this taxonomy is that it was collaboratively developed by a diverse group of scientists, including academic faculty, institutional leaders, and career development professionals. The diversity of this group and the group’s collective knowledge of job functions across sectors lends itself to the validity of this instrument. However, most were employed or have strong roots in academia, which likely affected taxonomy design and interpretation. Therefore, we must ask what biases were potentially introduced into the taxonomy due to the academic background of the taxonomy developers? To ensure inclusive representation from all career fields, we recommend the inclusion of employers in sectors outside the academy in subsequent discussions and improvements to UCOT v.2018.

One criticism of the current taxonomic nomenclature is the use of “Not Related to Science” and “Science-Related” in the classification of Career Type (*analogous to “Not Discipline Related”, as discussed above*), due to the negative connotation associated with a career that is not directly related to one’s training. Specifically, jobs categorized as “Not Related to Science” may be interpreted as unattractive, unusual, or not requiring or valuing PhD training. A similar consequence could stem from be categorized as “Other” *(for a discussion of the socially constructed “marked” vs. “unmarked” categories developed in relation to gender, race, etc., see* 31]). This taxonomy’s definition of “Not Related to Science” is “a career that is not directly relevant to the conduct of scientific research”, and, while we may agree that there are job types that fit that definition, one should not assume that the scientific skills and/or knowledge developed during research training are not used in those roles. For example, “children’s book author” may seem on the surface to be not related to science, however the author may publish books with science themes. Because detailed information on the nature of work will not be available in all cases, we accept that coders will need to make educated guesses for classification between "Not Related to Science" and other Career Types. We hope that, as scientists innovate new roles for themselves throughout the workforce, the predominant negative valuation of careers ‘not related to science’ will be challenged. We underscore that future iterations of this taxonomy could explore better ways to identify and classify the diverse career trajectories of PhD alumni without using “othered” terminology.

Finally, it has not escaped our attention that one shortcoming of all versions of this taxonomy— which was discussed at length during its generation and refinement over the past eighteen months— is its inability to sufficiently represent situations where an individual balances multiple job functions. For example, there is currently no way to notate the balance of research and teaching that is expected of faculty members who might otherwise be categorized as Career Type “Primarily Teaching” and Job Function “Teaching Faculty or Staff.” Physician scientists and other cross-functional roles provide the same challenge to accurate coding of their position. One potential solution for faculty roles involves elaborating the Faculty Flag fields to include additional yes/no drop-down prompts that investigate the contractual expectations of their appointment, e.g., research, teaching, clinical service, and/or service expectation. We recommend that the outcomes tracking community consider whether and how to address this currently unanswered problem.

## FUTURE DIRECTIONS

Although the evolving workforce will necessitate periodic updates to any classification scheme, we believe that broad adoption of UCOT v.2018 will aid institutions that have yet to adopt a taxonomy to categorize PhD career outcomes. Additionally, we recommend that a consortium of institutions using UCOT v.2018 be formed as soon as possible, with the purpose of convening users annually for the next 5 years to identify concerns about, or recommend modifications to, UCOT v.2018. To ensure that diverse perspectives and needs are considered, future meetings should include a collaborative cross-section of universities, workforce agencies, employers, and consortia, similar to the meeting convened by RBR to generate UCOT v.2017. We recommend that, as the taxonomy evolves, its reliability is periodically revisited and reported. Therefore, sustaining a unified taxonomy across many institutions will require working together as a community in an ongoing fashion.

Future studies to improve accuracy of career outcomes reporting could include triangulating classifications using multiple data sources, such as surveys of alumni and their employers, to confirm an administrative categorization of an individual’s Workforce Sector, Career Type, and Job Function. Alumni surveys might ask an individual to self-code their occupation, which could then be compared to the institutional classification. However, previous work from members of our working group (*GCM*) and others cast concerns about how reliably an alumnus/a could self-categorize their job without training in the nuances of a taxonomy. Therefore, using alumni for validation of their career categorizations (*via surveys, interviews, etc.*) warrants further thorough investigation. If institutions choose to include these time-intensive processes, we recommend that multiple university stakeholders (*e.g., registrar, human resources, graduate program, etc.*) synergize their priorities and streamline the internal processes to most effectively obtain as much information as possible from alumni. Finally, we suggest that universities, to whatever degree possible, build and maintain databases of contact information simultaneously and centrally with the collection and coding of career outcomes of its alumni.

As the academic community begins to embrace this important initiative of publicly sharing career outcomes, we recognize that institutions face difficult decisions when deciding how to prioritize competing demands on their budgetary and human capital resources. Therefore, for those institutions that have not yet begun classifying their alumni career outcomes, we recommend adopting the clarifications incorporated into UCOT v.2018 and its associated Guidance Document; we anticipate that the UCOT v.2018 will not only improve efficiency in coding, but also assure stakeholders that the data representing alumni career trajectories will be meaningful and consistent. We hope that UCOT v.2018 becomes a useful tool for institutions that are interested in providing transparency of career outcomes in ways that meet the practical needs of their graduate students, postdoctoral scholars, funding agencies, and other stakeholders.

The value of the Unified Career Outcomes Taxonomy v.2018 lies in its refined definitions and proscriptive Guidance Document, which together result in more reliable interpretation of the taxonomic categories. Applied correctly, it will impart consistency in publicly disseminated career outcomes data, ultimately empowering prospective and current graduate students and postdoctoral scholars to better understand the longitudinal career outcomes of trainees at individual institutions, and to make informed decisions about these prospective training environments.

## AUTHOR CONTRIBUTIONS

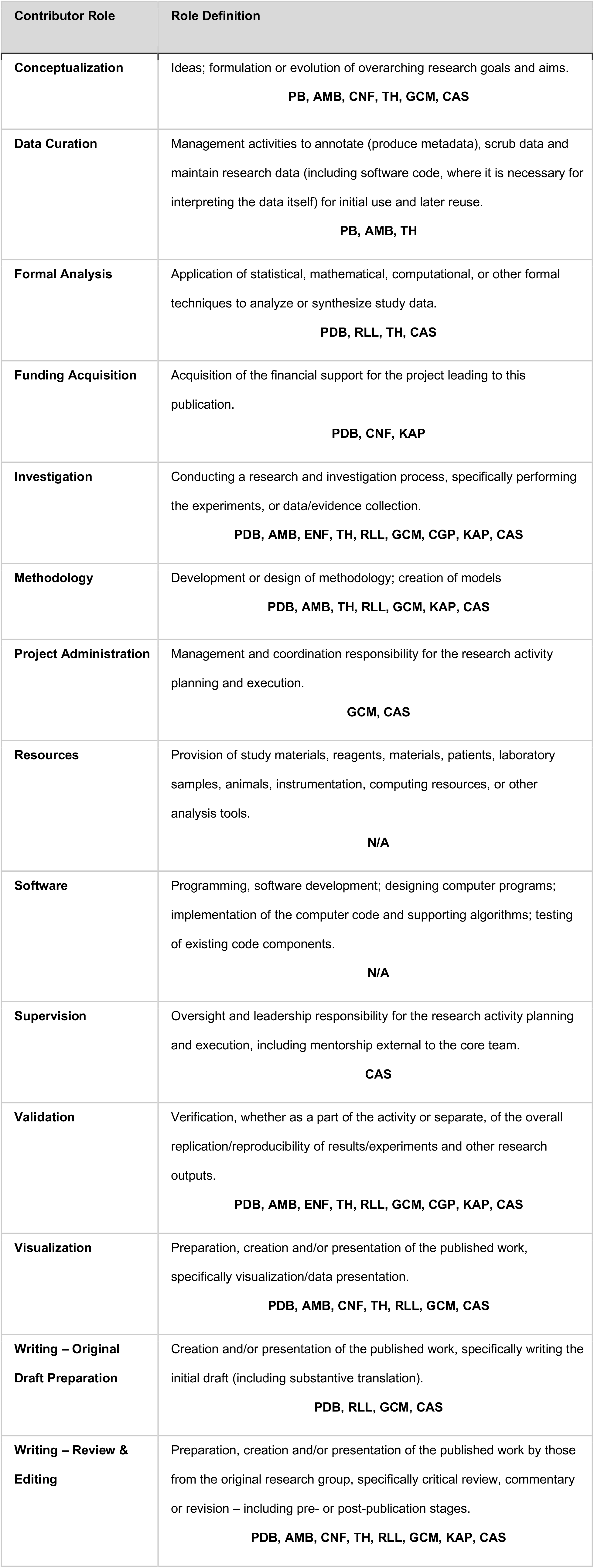

## ACKNOWLEDGEMENTS

We would like to thank members of the BEST Consortium, especially Julia Melkers, Liz Silva, Bill Lindstaedt, and Spencer Fenn for their valuable feedback on experimental design, data interpretation, and comments on earlier versions of this manuscript. We are grateful to Ambika Mathur, Roger Chalkley, and Fred Meyers for their foundational work in this field and continued support of this project. We thank Patricia Labosky, Rebecca Lenzi, and members of the BEST External Scientific Panel for their advocacy of this initiative. We thank the Odum Institute for Research in Social Science at the University of North Carolina at Chapel Hill for consultations on our statistical analyses, Brandon Barnes (Emory University) for his early contributions to data collection and curation methodology, and the Center for Postdoctoral Affairs in the Health Sciences at University of Pittsburgh for their thoughtful comments. The conclusions, views, and opinions expressed in this study are those of the authors and do not necessarily reflect the official policy or position of NIH, or the BEST consortium as a whole. The authors declare no competing interests.

## SUPPLEMENTAL MATERIALS

S1. Unified Career Outcomes Taxonomy v.2018 (unabridged)

S2. Annotated table of changes made to UCOT v.2017 in generating UCOT v.2018.

S3. Unified Career Outcomes Taxonomy v.2018 Guidance Document

S4. Selection and Composition of Alumni Records for Round 1 Data Set

S5. Selection and Composition of Alumni Records for Round 2 Data Set

S6. Two measures of discordance used to compare coding after Round 1 and 2

S7. Data Collection Workbook template.

